# Identification of novel myelodysplastic syndromes prognostic subgroups by integration of inflammation, cell-type composition, and immune signatures in the bone marrow

**DOI:** 10.1101/2024.03.11.584361

**Authors:** Sila Gerlevik, Nogayhan Seymen, Shan Hama, Warisha Mumtaz, I. Richard Thompson, Seyed R. Jalili, Deniz E. Kaya, Alfredo Iacoangeli, Andrea Pellagatti, Jacqueline Boultwood, Giorgio Napolitani, Ghulam J. Mufti, Mohammad M. Karimi

**Affiliations:** Comprehensive Cancer Centre, School of Cancer & Pharmaceutical Sciences, Faculty of Life Sciences & Medicine, King’s College London, London SE5 8AF, United Kingdom; Department of Basic and Clinical Neuroscience, King’s College London, London, United Kingdom; Department of Biostatistics and Health Informatics, King’s College London, London, United Kingdom; NIHR BRC SLAM NHS Foundation Trust, London, United Kingdom; Perron Institute for Neurological and Translational Science, University of Western Australia Medical School, Perth, WA 6009, Australia; Nuffield Division of Clinical Laboratory Sciences, Radcliffe Department of Medicine, University of Oxford, Oxford, United Kingdom

**Author notes:** These authors contributed equally to this work and share first authorship (Sila Gerlevik, Nogayhan Seymen, Shan Hama). The authors declare no competing financial interests.

## Abstract

Mutational profiles of Myelodysplastic syndromes (MDS) have established that a relatively small number of genetic aberrations, including SF3B1 and SRSF2 spliceosome mutations, lead to specific phenotypes and prognostic subgrouping. We performed a Multi-Omics Factor Analysis (MOFA) on two published MDS cohorts of bone marrow mononuclear cells (BMMNCs) and CD34+ cells with three data modalities (clinical, genotype, and transcriptomics). Seven different views, including immune profile, inflammation/aging, Retrotransposon (RTE) expression, and cell-type composition, were derived from these modalities to identify the latent factors with significant impact on MDS prognosis. SF3B1 was the only mutation among 13 mutations in the BMMNC cohort, indicating a significant association with high inflammation. This trend was also observed to a lesser extent in the CD34+ cohort. Interestingly, the MOFA factor representing the inflammation shows a good prognosis for MDS patients with high inflammation. In contrast, SRSF2 mutant cases show a granulocyte-monocyte progenitor (GMP) pattern and high levels of senescence, immunosenescence, and malignant myeloid cells, consistent with their poor prognosis. Furthermore, MOFA identified RTE expression as a risk factor for MDS. This work elucidates the efficacy of our integrative approach to assess the MDS risk that goes beyond all the scoring systems described thus far for MDS.

## Introduction

Myelodysplastic syndromes (MDS) are haematological diseases characterised by clonal proliferation due to genetic and epigenetic alterations within haematopoietic stem and progenitor cells (1, 2). Thus far, the prediction of a patient’s overall survival (OS) and event-free survival (EFS) has been predominantly dependent on the degree of peripheral blood cytopenias, bone marrow (BM) blast percentage, cytogenetics, and genetic features (3–6). This may, however, overlook other biological phenotypes and risk factors in MDS, including inflammation, age and aging characteristics, immune profile, BM cell-type composition, and expression of the noncoding sequences in particular retrotransposable elements (RTEs).

Inflammation can either suppress or promote cancer development (7). Inflammation can be protective against malignancies, including MDS, through innate and adaptive immune responses, enhancing antitumor immunity by promoting the maturation and function of dendritic cells (DCs) and initiating effector T cell responses (8). However, low-level chronic inflammation can result in an immunosuppressive milieu preventing the innate and T-cell antitumor immunity (9). Inflammaging is the process by which an age-related increase in chronic inflammation occurs, but the extent to which this process and other age-related events can impact the overall prognosis in MDS is yet to be uncovered (10, 11).

RTEs are genomic remnants of ancient DNA sequences that comprise a large part of the human non-coding genome and are evolutionarily silenced. RTEs include three classes: long terminal repeat (LTR) RTEs, and long and short interspersed nuclear elements, known as LINEs and SINEs, respectively. As part of the host defence mechanism, RTEs are silenced through DNA methylation and histone modifications in somatic cells (12). Mutations in genes implicated in DNA methylation and histone modifications (DNMT3A, TET2, ASXL1, and IDH1/2) are frequently reported in MDS. It is postulated that global hypomethylation can reactivate RTEs in MDS cases with epigenetic mutations, particularly in DNMT3A mutant cases, as has previously been shown for other cancer types (13, 14). Despite the availability of multiple transcriptomics data for MDS (15–18), a comprehensive assessment of RTE expression and its relationship to genetic variations or prognosis has not been documented.

Splicing factor (SF3B1 SRSF2, and U2AF1) mutations are the most common mutations in MDS (19–22). SF3B1 mutations are associated with the MDS ring sideroblasts type, good prognosis, and low leukemic transformation (20). In contrast, SRSF2 mutations are associated with poor prognosis and are more prevalent in the male sex and older age (23). Recent studies have shown mutations in splicing factors can induce chronic innate immunity and enhance NF-κB signalling in MDS through aberrant splicing of various target genes. (18, 24, 25). These recent mechanistic studies have significantly advanced our understanding of the consequences of splicing factor mutation in human and model organisms. However, we still lack a systematic approach to integrate splicing factor mutations with other players in the tumour microenvironment, including immune profile and BM cell-type composition.

Recent work has explored the relationship between transcriptional signatures and critical signalling pathways to determine survival prognosis and diagnostic efficacy in MDS patient cohorts (26). However, no studies to date integrated clinical MDS phenotypes, RTE expression, cell-type composition, and immune and aging gene signatures. This study employs MOFA for a comprehensive analysis of three data modalities (clinical, genotypic, and transcriptomic) and seven different “views” derived from these modalities to identify the factors that may impact MDS prognosis. MOFA could not identify any factor representing splicing factor mutations; hence, we examined our entire feature sets from cell-type composition, immune profile, and inflammation/aging views to identify the features associated with mutations in SF3B1 and SRSF2 genes in MDS cohorts.

## Methods

### MDS RNA-seq cohorts

We utilised two RNA-seq datasets for MDS (**Supplementary Table 1**). The first was data from bone marrow mononuclear cells (BMMNCs) of 94 MDS patients obtained from the Shiozawa *et al.* study (17) that is enriched with splicing factor mutations (**Supplementary Table 2**). The second dataset utilised bone marrow CD34+ haematopoietic stem and progenitor cells (HSPCs) data that Pellagatti *et al.* derived from 82 MDS patients (15), which again focused on MDS cases with splicing factor mutations (**Supplementary Table 2**).

### Generating gene signature scores using *singscore*

For the immunology and inflammation/aging gene sets, we performed a meticulous literature review and generated a list of gene sets from previously published articles (**Supplementary Table 3**). The markers for the cellular composition gene sets were taken from van Galen *et al.* (27). For each of the curated gene sets, instead of looking at individual gene expressions, we used *singscore* (version 1.20.0) (28), a method that scores gene signatures in a cohort of samples using rank-based statistics on their gene expression profiles. For RNA-seq data, we provided Reads per Million (RPM) normalized expression values to *singscore*. In the case of microarray data, we first compiled the microarray expression matrix for average expression values for all probes overlapping each gene using the *limma* package (29) in R; then we used the rankGenes function from the *singscore* package to rank each gene sample-wise. Eventually, the multiScore function was used to calculate signature scores for all gene sets at once.

### Multi-Omics Factor Analysis (MOFA)

MOFA was applied on seven views derived from the BMMNC and CD34+ cohorts: immune profile, cell-type composition, inflammation/aging, genotype, RTE expression, clinical numeric, and clinical categorical views, using the *MOFA2* package (version 1.10.0) in R (**Fig. 1a**). Each view consists of non-overlapping features of the same sample set of patients. Since aging and inflammation views share some gene sets, including inflammatory chemokines and cytokines, and we did not want to repeat these features through MOFA analysis, we combined these two views as the inflammation/aging views throughout this study. The views were scaled to have the same unit variance via the scale_view option from the MOFA model. The model pruned inactive factors incapable of capturing significant variance within the biological views, generating ten factors with minimum explained variance 2% in at least one biological view. Significant features within each view were determined based on the absolute weight threshold above 0.5 in at least one of the ten identified factors.

**Figure 1.**
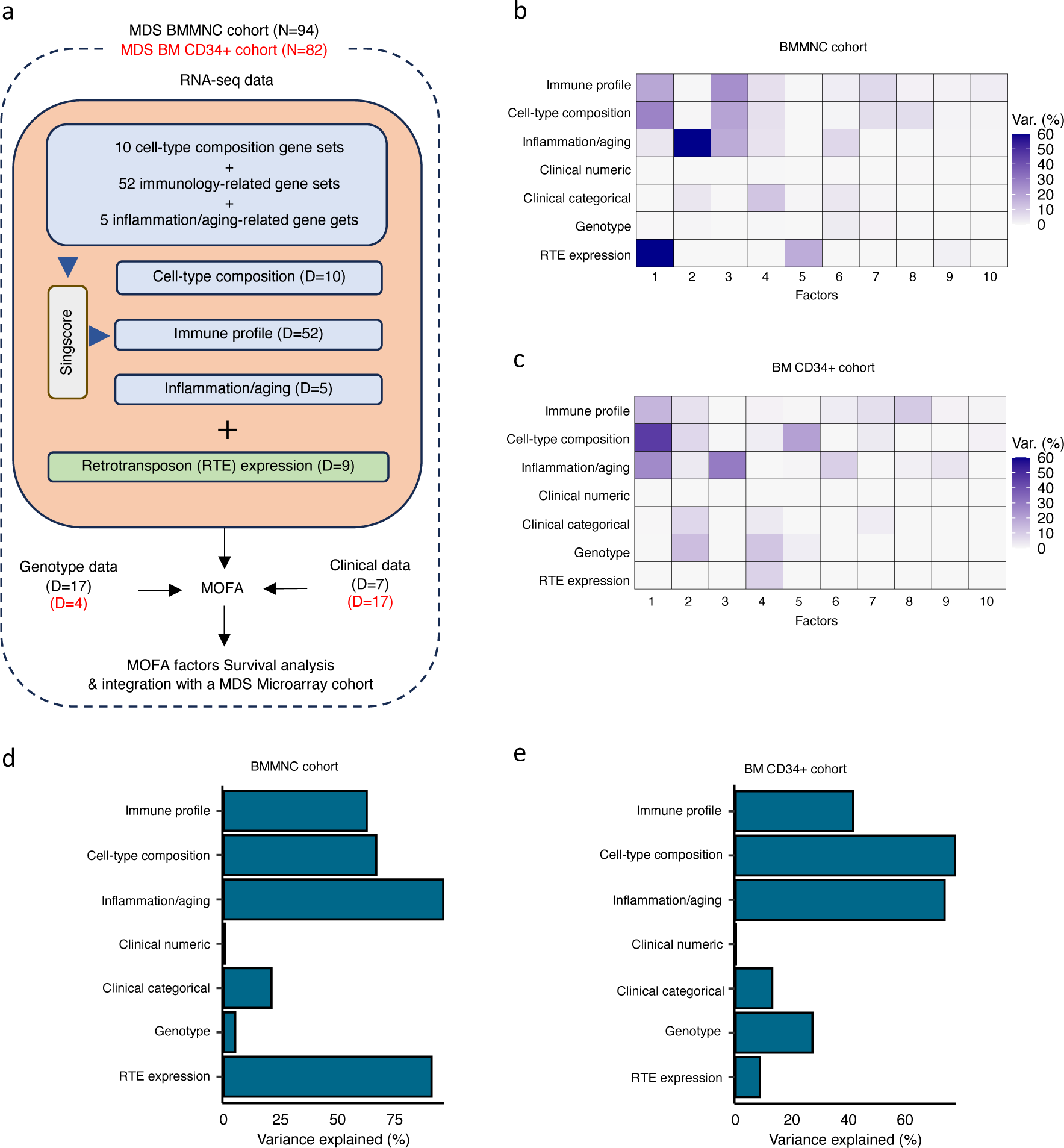
Schematic of MOFA workflow overview, downstream analyses, and factor determination in the BMMNC and BM CD34+ cohorts. **a,** RNA-seq, genotype and clinical data were obtained from bone marrow mononuclear cell (BMMNC) samples of 94 MDS patients from Shiozawa *et al.* and BM CD34+ samples of 82 patients from Pellagatti *et al.* studies. We generated seven views of the data where three of which were derived from RNA-seq data after applying *Singscore*: immune profile, cell-type composition, inflammation/aging. The other four views were clinical numeric and categorical (**Supplementary Table 4**), genotype and retrotransposable element (RTE) expression. The data were put through MOFA to identify latent factors and the variance decomposition by factors. The number of features (dimensions) per view is abbreviated by “D”. **b-c,** The determined factors for the BMMNC and BM CD34+ cohorts and the percentage of explained variance for each view per identified factor were shown. **d-e,** Bar charts depict the total variance explained for each biological data view by all the factors combined in the BMMNC and BM CD34+ cohorts.

### Survival analysis

The association between latent factors and survival outcomes was investigated with Cox regression analysis and Kaplan-Meier curves via the R package *survival* (version 3.5-5). Within the BMMNC cohort, overall and event-free survivals were used as separate response variables in the univariate Cox regression, with latent factors employed as predictors. Additionally, in the multivariate Cox regression, age and sex were included as predictors alongside the factors. In the CD34+ cohort, only overall survival was used as the response variable for both regression analyses. Kaplan-Meier plots were constructed for factors exhibiting a significant hazard ratio in the univariate Cox regression, categorizing factor values into three groups: ‘low’ for the 1st quartile, ‘high’ for the 4^th^ quartile, and ‘intermediate’ otherwise. The statistical significance between the high and low factor groups was determined using the log-rank test via the R package *survminer* (version 0.4.9).

### Differential Expression and Gene Set Enrichment Analysis (GSEA)

We performed differential expression analysis in high versus low Factor 1 groups using *DESeq2* (30) and generated p-values and statistics for each gene. The genes were sorted based on the “stat” column in *DESeq2* and provided to the GSEA software (31) as an input. GSEA was separately run on cancer hallmark and Reactome gene set databases. The GSEA output was the list of up or down-regulated pathways from each database.

### RTE expression

To generate the RTE expression, we mapped the RNA-seq reads to RepeatMasker to extract the reads covering the RTE regions and calculated the Reads Per Million (RPM) scores for each class and family of RTEs. We included nine families from three main RTE classes: 1) CR1, L1 and L2 families from LINE; 2) Alu and MIR from SINE; and 3) ERV1, ERVL, ERVL-MaLR, and ERVK families from LTR.

## Results

### MOFA identified latent factors linking different views in multimodal MDS data

We applied MOFA to identify latent factors within BMMNC and CD34+ MDS cohorts (**Fig. 1a**). To run MOFA on these two MDS cohorts, three (immune profile, inflammation/aging profile, and cell-type composition) out of seven views were derived from RNA-seq by applying *singscore* (28) on RNA-seq gene expression. Each of these three views were carried forward in the workflow by a number of gene sets, and per gene set, *singscore* generated relative gene signature scores for all samples within each cohort (**Fig. 1a** and **Supplementary Table 3**). The other four views were clinical numeric, clinical categorical, genotype, and RTE expression.

We have provided the data generated for these seven views to MOFA in two separate runs for BMMNC and CD34+ MDS cohorts. For each cohort, we could identify ten factors (minimum explained variance 2% in at least one biological view) from the 15 default factors generated by MOFA (**Fig. 1b-c**). These factors displayed a relationship with different biological views, with Factor 1 as the most dominant factor, linking immune profile, cell-type composition, and inflammation/aging profile in both cohorts. Further to this, in the BMMNC cohort, a high level of variance for RTE expression was explained by Factor 1 (**Fig. 1b**). We also observed similar trends in terms of the level of variance explained by the identified factors for the biological views in both cohorts, with the only notable difference being a greater level of variance explained for genotype data and lower level of variance explained for RTE expression in the BM CD34+ cohort versus the BMMNC cohort (**Fig. 1d-e**). Additionally, dividing the patients based on low (1^st^ quartile), intermediate (2^nd^ and 3^rd^ quartile), and high (4^th^ quartile) levels of Factor 1 in both cohorts could successfully stratify patients in the dimensionality reduction plots obtained from applying principal component analysis (PCA) on gene expression data from both cohorts (**Supplementary Fig. 3**).

MOFA also identified the highly weighted features within each factor in the BMMNC and BM CD34+ cohorts (**Fig. 2a-b**), in which each feature belongs to a specific biological view. We further characterised Factor 1 as the most dominant factor linking multiple features from different views. We observed that high Factor 1 in the BMMNC cohort represented patients who have cells with stem and progenitor-like characteristics. This includes but is not limited to a positive correlation with features comprising progenitor-like and HSC-like (**Fig. 2c**). In contrast, there is an inverse correlation between GMP/GMP-like and Factor 1 scores (**Supplementary Fig. 1c**). SINE:Alu expression is increased in patients high in Factor 1 within this cohort, representing a group that may have significant levels of genetic instability (**Supplementary Fig. 1a**). Moreover, Factor 1 correlates with the following immunology features: increased T-helper 1 (Th1) cells, and decrease in certain immune cells, especially in neutrophils and exhausted CD8+ T cells (**Supplementary Fig. 1b**). The relationship between Th1 and SINE elements is particularly interesting given the role of Th1 cell activation in host defence against inflammation that may be modulated by SINE activation. Furthermore, there is a modest correlation between immunosenescence and exhausted CD8+ T cells scores, particularly in patients with higher levels of GMPs, demonstrating the ineffectiveness of the immune system in GMP-dominant patients (**Fig. 2c**).

**Figure 2.**
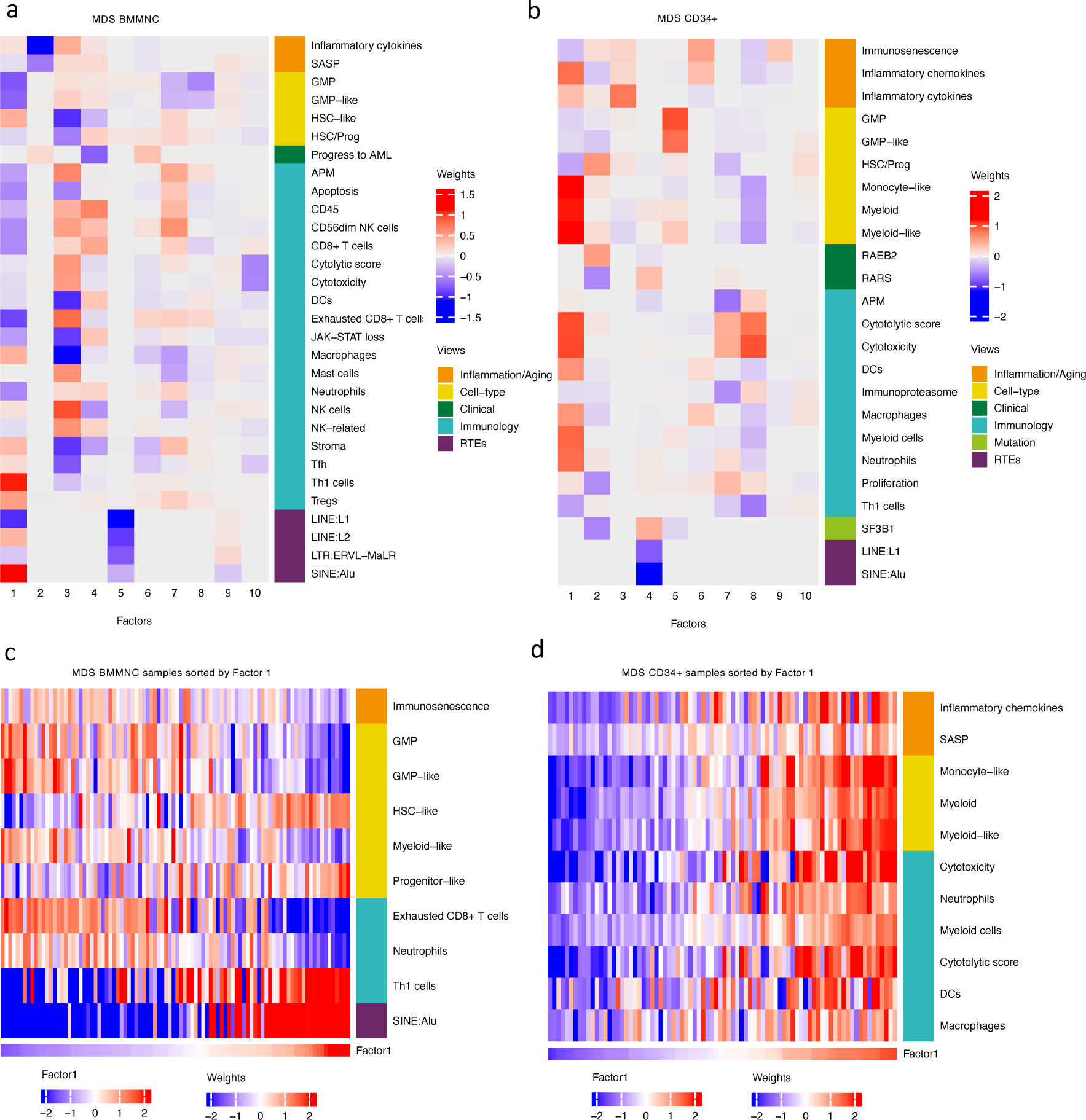
Breakdown of important features for each factor generated by MOFA. **a-b,** The important features with high weights for each biological view per factor were shown for the BMMNC and BM CD34+. Blue represents features with the inverse correlation with the factor, and red shows the positive correlation. **c-d,** Characterisation of Factor 1 in the BMMNC and BM CD34+ cohorts, showing only those features highly influencing Factor 1 for the patients in these cohorts. Patients were sorted by Factor 1 values.

MOFA analysis of the CD34+ MDS Cohort revealed factors associated with different immune signatures. Factor 1 is associated with signatures of differentiated myeloid cells (**Fig. 2d** and **Supplementary Fig. 2c**), Factor 2 with HSPC, and Factor 4 with GMP progenitor cells, suggesting that these factors might correlate with distinct differentiation states of MDS blasts (**Fig. 2b**). Factor 1 is also associated with cytolytic and cytotoxicity score (**Fig. 2d** and **Supplementary Fig. 2b**). Since CD34+ cells used for this analysis were purified using anti-CD34 conjugated beads and not FACS sorted, this signal might depend on the presence of T or NK cells contaminations in the CD34+ fraction. The trend that we observed for Factor 1 prompted us to conduct a differential expression and gene set enrichment analysis (GSEA) to identify the pathways that are dysregulated in patients having high (4th quartile) versus low (1st quartile) levels of Factor 1. Interestingly, we found upregulation of inflammatory, Interferons, TNFA, JAK-STAT signalling pathways within cancer hallmark gene sets **(Supplementary Fig. 4a)** and upregulation of chemokines and Neutrophils amongst Reactome gene sets **(Supplementary Fig. 4b)** supporting the trend observed by MOFA (**Fig. 2d** and **Supplementary Fig. 2b**). This analysis suggests Factor 1 as an immune-active factor associated with high-level of cytotoxicity and inflammation in the CD34+ MDS Cohort.

### The Cox regression models identified the high expression of RTEs as a risk factor and inflammation as a protective factor in MDS

We explored the power of the latent factors generated by MOFA as predictors of MDS survival. Univariate and multivariate Cox regression models identified three of the ten factors identified by MOFA were significantly associated with OS or EFS in the BMMNC cohort (**Table 1**). The univariate analysis was conducted to investigate the association of the factors with survival, and the multivariate analysis controlled for sex and age. Our univariate analysis demonstrated that Factors 4 and 9 significantly impact OS (HR: 0.39; P = 0.028 and HR: 0.49; P = 0.012, respectively) and EFS (HR: 0.48; P < 0.001 and HR: 0.59; P = 0.007, respectively) in patients in the BMMNC cohort (**Table 1**). Interestingly, the impact of Factor 4 on EFS became insignificant after controlling for sex and age. Though no significance was associated with OS and Factor 2, EFS in both the univariate and multivariate analyses exhibited a statistically significant association with this factor (HR: 1.7; P = 0.033 and HR: 2.27; P = 0.033, respectively) (**Table 1**). The 10 factors identified by MOFA in BM CD34+ cohort did not show any significance associated with MDS overall survival (**Supplementary Table 5**).

Dividing the patients based on low (1^st^ quartile), intermediate (2^nd^ and 3^rd^ quartile), and high (4^th^ quartile) levels of Factors 2, 4, and 9 in the BMMNC cohort, we found Factors 4 and 9 exert a protective influence over the prognosis of MDS patients (**Supplementary Fig. 5a** and **Fig. 3a**, respectively), whilst higher levels of Factor 2 predict a poorer prognosis for these same patients (**Fig. 4a**). Upon investigation of the RTE absolute loading of features in Factor 9, patients with low Factor 9 level showed increased LTR:ERV1, SINE:MIR and SINE:Alu demonstrating the high level of RTE expression as a potential risk factor for MDS (**Fig. 3a-b**). In contrast, for Factor 2, some component weights for inflammation/aging including inflammatory cytokines and SASP were increased once the Factor 2 level is decreased suggesting the secretion of cytokines and the down-stream inflammation as a protective factor for MDS (**Fig. 4a-b**).

**Figure 3.**
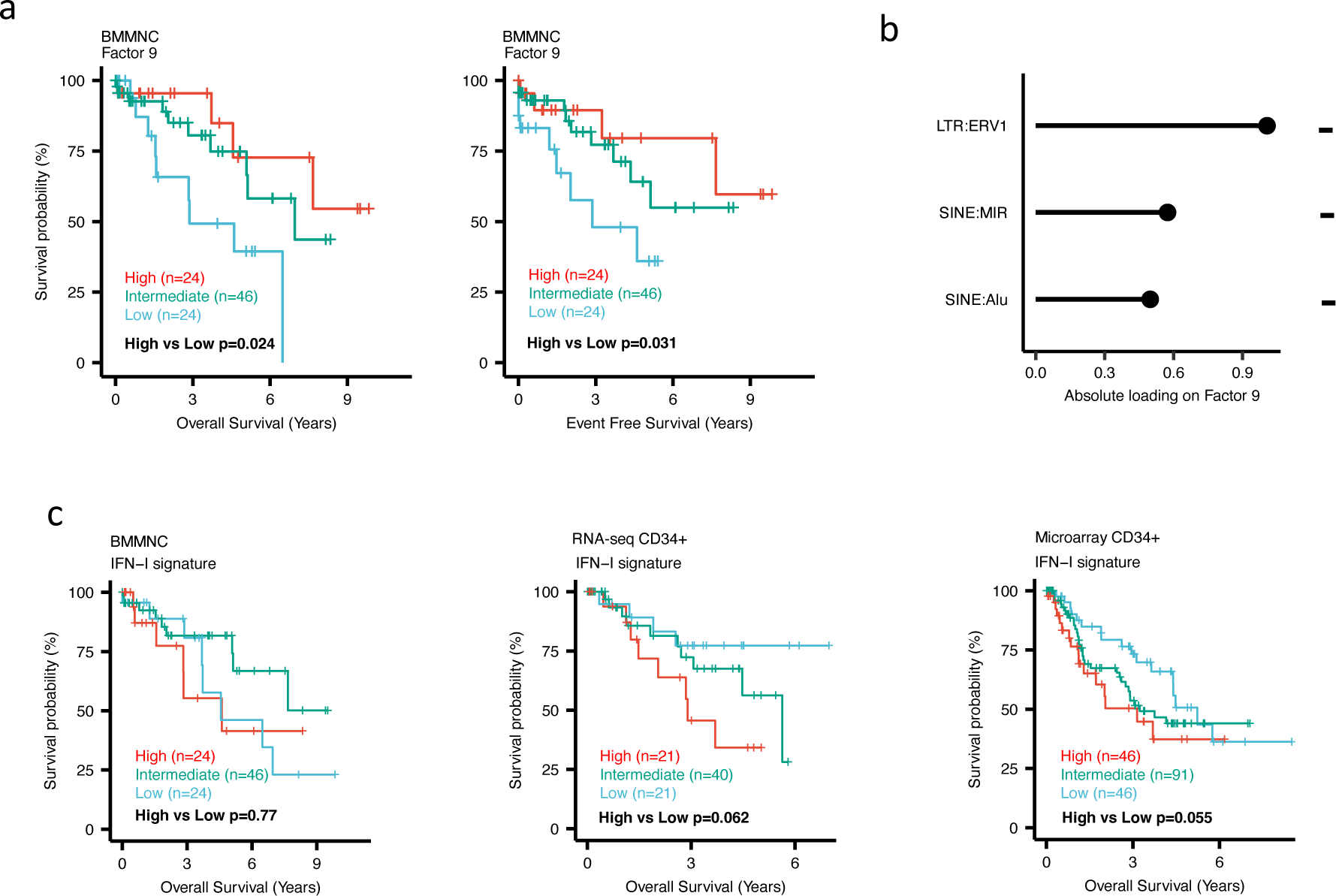
Impact of Factors 9 and IFN-I levels on MDS prognosis. **a,** Kaplan-Meier plots for the BMMNC cohort where patients were split based on low (1^st^ quartile), intermediate (2^nd^ and 3^rd^ quartile), and high (4^th^ quartile) levels of Factor 9. **b,** The absolute loading of the top three features affecting Factor 9 in RTE expression view in the BMMNC cohort. **c,** Kaplan-Meier plots where patients were split into quartiles (high 25%, low 25% and intermediate 50%) for survival analyses depending on their IFN-I signature score levels in the BMMNC, RNA-seq CD34+, and Microarray CD34+ cohorts. All p-values were calculated using log-rank test on overall and event-free survival values of high versus low groups.

**Figure 4.**
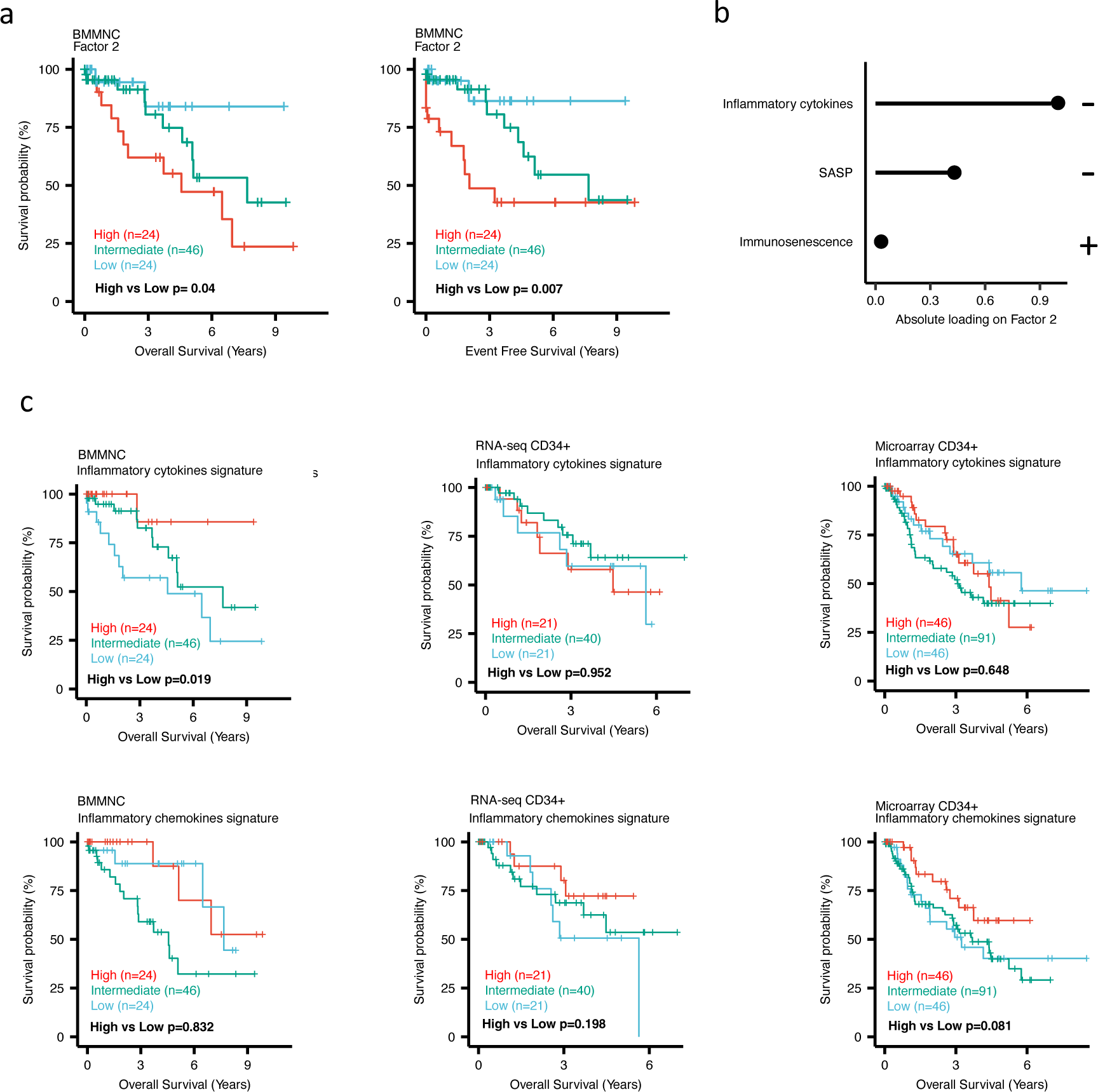
Impact of Factors 2 and inflammation levels on MDS prognosis. **a,** Kaplan-Meier plots for the BMMNC cohort where patients were split based on low (1^st^ quartile), intermediate (2^nd^ and 3^rd^ quartile), and high (4^th^ quartile) levels of Factor 2. **b,** The absolute loading of the top three features affecting Factor 2 in inflammation/aging biological view in the BMMNC cohort. **c,** Kaplan-Meier plots where patients were split into quartiles (high 25%, low 25% and intermediate 50%) for survival analyses depending on their inflammatory cytokines and chemokines levels in the BMMNC, RNA-seq CD34+, and Microarray CD34+ cohorts. All p-values were calculated using log-rank test on overall and event-free survival values of high versus low groups.

Factor 4 in the BMMNC cohort shows several data views and displays a more specific phenotype of the patients. This phenotype includes accumulation of immune cells, such as CD56dim NK cells, CD8+ T cells, DCs, and neutrophils (**Fig 2.a**). Thus, Factor 4 points to a more immune-active disease. In line with this, there is a decreased rate of the ‘progress to AML’ phenotype (**Fig 2.a** and **Supplementary Fig 5.b**) in patients with high level of Factor 4. In contrast, for the patients showing low level of Factor 4, we observed that the number of healthy stem and progenitor cells (HSCs) decreased (**Supplementary Fig 5.c),** whereas the number of malignant HSCs (HSC-like) increased (**Supplementary Fig 5.c-d)**. Most notably, the BM of these patients have a high content of stroma and a low content of leukocytes, represented by low CD45 marker score (**Supplementary Fig 5.c and e**). Overall, this result suggests that for a subset of MDS patients, represented by the low level of Factor 4, the haematopoiesis is impaired due to the depletion of healthy HSCs, leading to the decrease in the number of leukocytes, an increase of stroma, and features of secondary AML.

### BM CD34+ cohorts support better prognosis for high-inflamed cases but poor prognosis for IFN-1-induced cases

To better understand the drivers of prognosis for MDS, patients were split into quartiles (high 25%, low 25% and intermediate 50%) for survival analyses depending on their levels for individual features (inflammatory cytokines, inflammatory chemokines, and IFN-I signature) in the two MDS cohorts. In addition, a microarray gene expression dataset obtained from the HSPCs isolated from BM of 183 MDS patients (16) was also included in this analysis. Recall, in the BMMNC cohort, Factor 2 was primarily represented by the inflammation/aging biological view (**Fig. 4b**) and Factor 9 was represented by RTE expression (**Fig. 3b**). To confirm the isolated impact of the aforementioned features on patient survivals in CD34+ cohorts, we created survival plots for each of the three cohorts separating patients into quartiles based on high, low and intermediate levels of the features. The only situation in which we observed statistical significance was the high score of inflammatory cytokines in the BMMNC cohort, which was associated with superior OS (P = 0.019) (**Fig. 4c**) supporting the strong contribution of this feature to Factor 2 (**Fig. 4b**). For the rest of the analyses, statistical significance was not achieved, though there may be a trend pertaining to higher inflammatory chemokines signature and better prognosis in CD34+ cohorts (**Fig. 4c**). Further to this, higher IFN-I signature score was also related to a poorer prognosis in CD34+ cohorts (**Fig 3.c**). Strikingly, this pattern was not observed in the BMMNC cohort (**Fig 3.c**), suggesting an alternative mechanism other than IFN-I activation can contribute to poor MDS prognosis due to retroviral activations (**Fig 3.a-b**).

### SF3B1 mutant cases show low HSPC content and high levels of inflammation in the BMMNC and BM CD34+ cohorts

When investigating the inflammation/aging and cell-type views in both the BMMNC and BM CD34+ cohorts, we identified a significant association between the occurrence of SF3B1 mutation and the high level of inflammatory chemokines (BMMNC P < 0.01; BM CD34+ P < 0.05) (**Fig. 5a-c**). However, we needed to know whether the co-occurrence of other mutations with SF3B1, as confounding factors, can affect the statistical significance of the association of SF3B1 mutation with inflammatory chemokines. Therefore, we applied a multiple linear regression model to examine the association between various mutations (covariates) and inflammatory chemokines while controlling the effect of other mutations. Strikingly, out of 13 different mutations in the BBMNC cohort, SF3B1 mutation was the only mutation showing a significant association with the inflammatory chemokine level in the multiple linear regression model (P < 0.0001) (**Table 2**).

**Figure 5.**
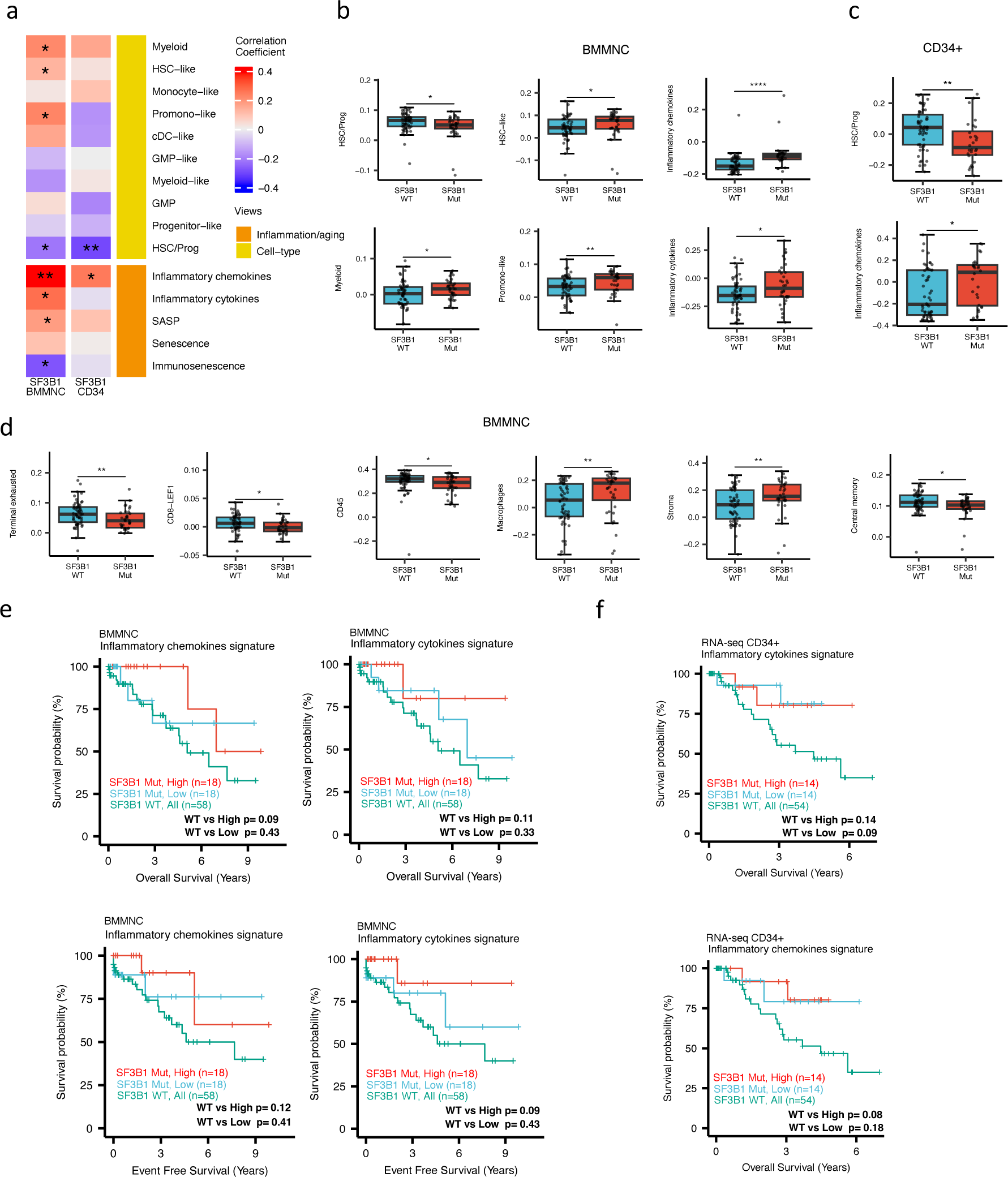
Characterisation of SF3B1 mutant MDS using gene signatures from multiple biological views. **a,** Association of a subset of inflammation/aging and cell-type features with SF3B1 mutation in the BMMNC and BM CD34+ cohorts, with red depicting a positive correlation and blue an inverse correlation with SF3B1 mutation. The significances were calculated with the Wilcox rank-sum test, and the significant associations were shown by * (P < 0.05), ** (P < 0.01) or *** (P < 0.001). **b-c,** Boxplots comparing the levels of the significant individual features from the cell-type and inflammation/aging biological views for SF3B1 mutant versus SF3B1 WT cases in the BMMNC and CD34+ cohorts. **d,** Boxplots comparing the levels of the significant individual features from the immune profile biological views for SF3B1 mutant versus SF3B1 WT cases in the BMMNC cohort. **e-f,** Kaplan-Meier plots displaying overall and event-free survivals for SF3B1 mutant cases split by high and low levels of inflammatory cytokines and chemokines versus SF3B1 mutant WT in the BMMNC and CD34+ cohorts. Event-free survival data was only available for the BMMNC cohort. All p-values were calculated using log-rank test on overall and event-free survival values of SF3B1 mutant high and low versus WT groups.

In both cohorts, the SF3B1 mutant pertained to decreased HSC/Prog cells (BMMNC P < 0.05, BM CD34+ P < 0.01) (**Fig. 5a-b**). Furthermore, the BMMNC cohort saw a statistically significant increase in myeloid, HSC-like, and promo-like scores, along with inflammatory cytokines level (P < 0.05) in SF3B1 mutant cases (**Fig. 5a-b**). Applying multiple linear regression models and controlling the effect of other mutations supported the identified correlation between SF3B1 mutation and inflammatory cytokines level (P < 0.02) (**Table 2**). We then examined the Immunology view in the BMMNC cohort to see whether we could identify any immune signature that the SF3B1 mutation may influence (**Supplementary Fig. 6a**). We found that SF3B1 mutants tend to have low levels of leukocytes but high levels of stroma and macrophages (**Fig. 5d**). This result supports previous findings by Pollyea et al. (32, 33) demonstrating the generation of inflammatory cytokines due to macrophage activation in human BM samples obtained from SF3B1 mutant patients.

We also divided the SF3B1 mutant cases from the BMMNC cohort into high versus low groups depending on their levels of inflammatory chemokines and cytokines. Although survival plots did not reach statistical significance, the trend shows that SF3B1 mutant cases with higher inflammation tend to survive better than wild-type SF3B1 (**Fig. 5f-g**). This trend indicates that the SF3B1 mutation perhaps only induces macrophage activation to survive better in a subset of mutant cases.

### SRSF2 mutant cases show high GMP content and high levels of senescence and immunosenescence

When assessing inflammation/aging and cell-type features for MDS SRSF2 mutant cases in both cohorts, we saw a statistically significant increase in GMP and GMP-like features (P < 0.05), which are precursor cells to granulocytes and monocytes (**Fig. 6a-c**). The increased frequency of GMPs is an inherent feature of high risk MDS (34, 35). In the BMMNC cohort, within *SRSF2* mutant cases, there was a statistically significant decrease in healthy myeloid features (P < 0.05), along with an increase in myeloid-like (malignant myeloid) and senescence/immunosenescence (P < 0.05) (**Fig. 6a-b**). Further analyses of the BMMNC cohort indicated that SRSF2 mutants tend to have low levels of T cells and a reduction of T cell activity. The levels of several immune cells were decreased in SRSF2 mutants, including T, Th1, and Treg cells (P < 0.05) (**Fig. 6d and Supplementary Fig. 6b**). Patients with SRSF2 mutation, related to poorer prognosis in literature, also had a lower cytolytic score (P < 0.001) and increased central memory cells (P < 0.01) (**Fig. 6d)**. Taken together, this result finds relevance to the immunosenescence and the reduction of the T cells and their cytolytic activity in SRSF2 mutant MDS, leading to a severe and high-risk situation for MDS.

**Figure 6.**
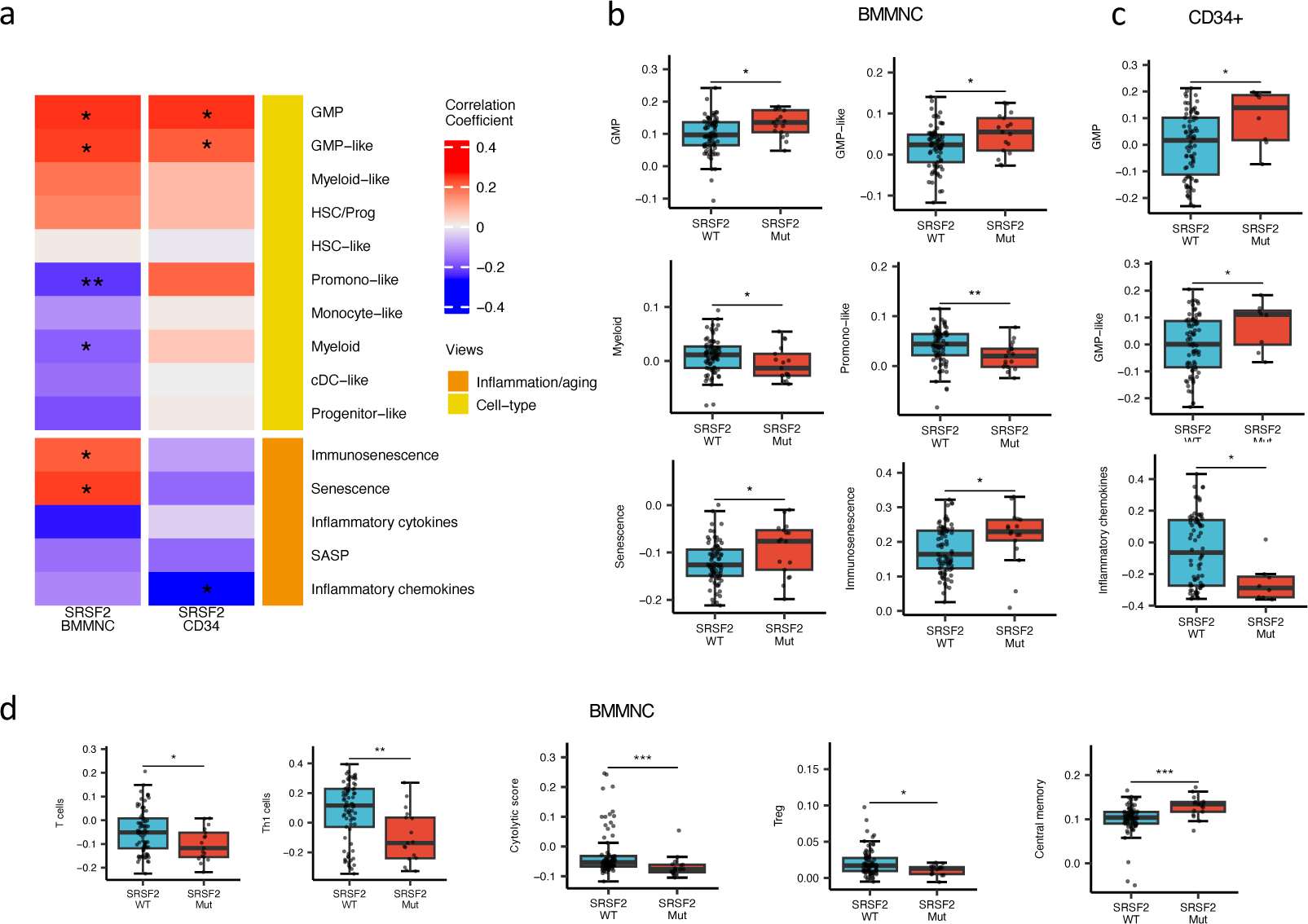
SRSF2 mutant MDS is catheterised by high GMP content and high levels of senescence and immunosenescence. **a,** Association of a subset of inflammation/aging and cell-type features with SRSF2 mutation in the BMMNC and BM CD34+ cohorts, with red depicting a positive correlation and blue an inverse correlation with SRSF2 mutation. The significances were calculated with the Wilcox rank-sum test, and the significant associations were shown by * (P < 0.05), ** (P < 0.01) or *** (P < 0.001). **b-c,** Boxplots comparing the levels of the significant individual features from the cell-type and inflammation/aging biological views for SRSF2 mutant versus SRSF2 WT cases in the BMMNC and CD34+ cohorts, respectively. **d,** Boxplots comparing the levels of the significant individual features from the immune profile biological views for SRSF2 mutant versus SRSF2 WT cases in the BMMNC cohort.

## Discussion

We applied MOFA to seven biological views derived from two MDS patient cohorts. MOFA could identify latent and important phenotypes from multimodal MDS data. Notably, we identified RTE expression as a risk factor and inflammation as a protective factor in MDS. Moreover, we uncovered that Factor 4 correlated with progress to AML, with patients lower in this factor more likely to develop AML. Low level of Factor 4 is related to a phenotype with low frequency of leukocytes; patients also typically had increased malignant HSCs and stroma. Our analysis shows that these patients probably have a defect in haematopoiesis, preventing the production of sufficient blood cells and ultimately allowing for the stroma to invade the whole marrow (36, 37). Although this observation is not novel, it is confirmatory of prior researches and demonstrates the power of our approach for integrating multimodal MDS data.

Literature has clearly defined the relationship between the SF3B1 mutation and long OS and EFS, with a low risk of progression to AML (23, 38, 39). We showed that SF3B1 mutant MDS cases tend to have high levels of inflammation, perhaps due to macrophage activation, and thus confers good prognosis for patients in terms of anti-tumour activity. Therefore, inflammation might help MDS survival. Further division of SF3B1 mutant patients into high and low levels of inflammation shows that the SFSB1 mutants with higher inflammation can generally survive better despite not being statistically significant. Investigating the synergic factors including epigenetic factors remains elusive. Enriching the multi-omics data with new modalities, including epigenomes, CyTOF, and Luminex cytokine/chemokine data in the future might help us understand why SF3B1 mutants show distinct patterns in terms of inflammation and survival.

Our work revealed that SRSF2 mutant MDS cases show a reduction of T cells. The decrease in the ability of patients to accumulate T cells, such as Th1 cells, which play crucial roles in modulating killing of tumour cells may cause worse outcomes for SRSF2 mutant patients (40).

We also observed increased immunosenescence level in SRSF2 mutants. Recent studies have shown that the expression of programmed death-ligand 1 (PD-L1) protein is significantly elevated in senescent cells (41–43). Increased PD-L1 protein levels protect senescent cells from being cleared by cytotoxic immune cells that express the PD-1 checkpoint receptor. In fact, activation of the PD-1 receptor inhibits the cytotoxic capabilities of CD8+ T and NK cells, increasing immunosenescence. Notably, patients with MDS who possess particular somatic mutations, such as those in the TP53, ASXL1, SETBP1, TET2, SRSF2, and RUNX1 genes, have an increased propensity to react favourably to PD-1/PD-L1 inhibitors (44) confirming that many cellular and molecular mechanisms, known to promote cellular senescence, including alteration of splicing machinery, are crucial stimulators of the expression of PD-L1 protein. Interestingly, in our analysis, we also observed a correlation between the senescence gene signature score and the expression of the PD-L1 gene in CD34+ cells (**Supplementary** Figure 7), supporting the previous findings linking PD-L1 gene expression to cellular senescence.

The immunology and ageing features extracted from the MDS transcriptomic data used in our analysis pipeline can enhance the conventional risk-scoring systems for MDS by providing new insights into this disease, particularly in the context of inflammation and ageing. For some patients, the clinical and genetic features may remain relatively the same until follow-up. Still, the transcriptomic features might differ considerably from the baseline diagnosis, affecting the course of treatment.

This study contributed to a deeper understanding of MDS pathogenesis and identified potential prognostic markers for this disease. It also elucidated the importance of considering the relationships between different pathways, markers, and mutations in predicting patient outcomes, highlighting the efficacy of a comprehensive approach that goes beyond all the scoring systems that have been described thus far for MDS.

## Supporting information

Manuscript Tables

## Acknowledgment

This study was supported with funding from Bristol Myers Squibb (BMS) company during this project. We thank Dr Sh. Kordasti for critical reading of the manuscript. AP and JB were supported by Blood Cancer UK (grants 13042 and 19004).

## Authors’ contributions

MMK and GJM conceived this analysis. The analysis was mainly performed by SG, and NS with contributions from WM, IRT, and DEK. MMK, SG, and NS designed the graphics. MMK, SG, SH, NS, and SRJ wrote the manuscript with input from all authors. GN, SO, AP, and JB contributed to the interpretation of the analytics. AL provided statistical support during the study. MMK and GJM supervised the work. All authors read and approved the final manuscript.

## Data Availability

Scripts and data used in this study are available on Github (https://github.com/Karimi-Lab/MDS_MOFA).

---

**Supplementary Figure 1.**
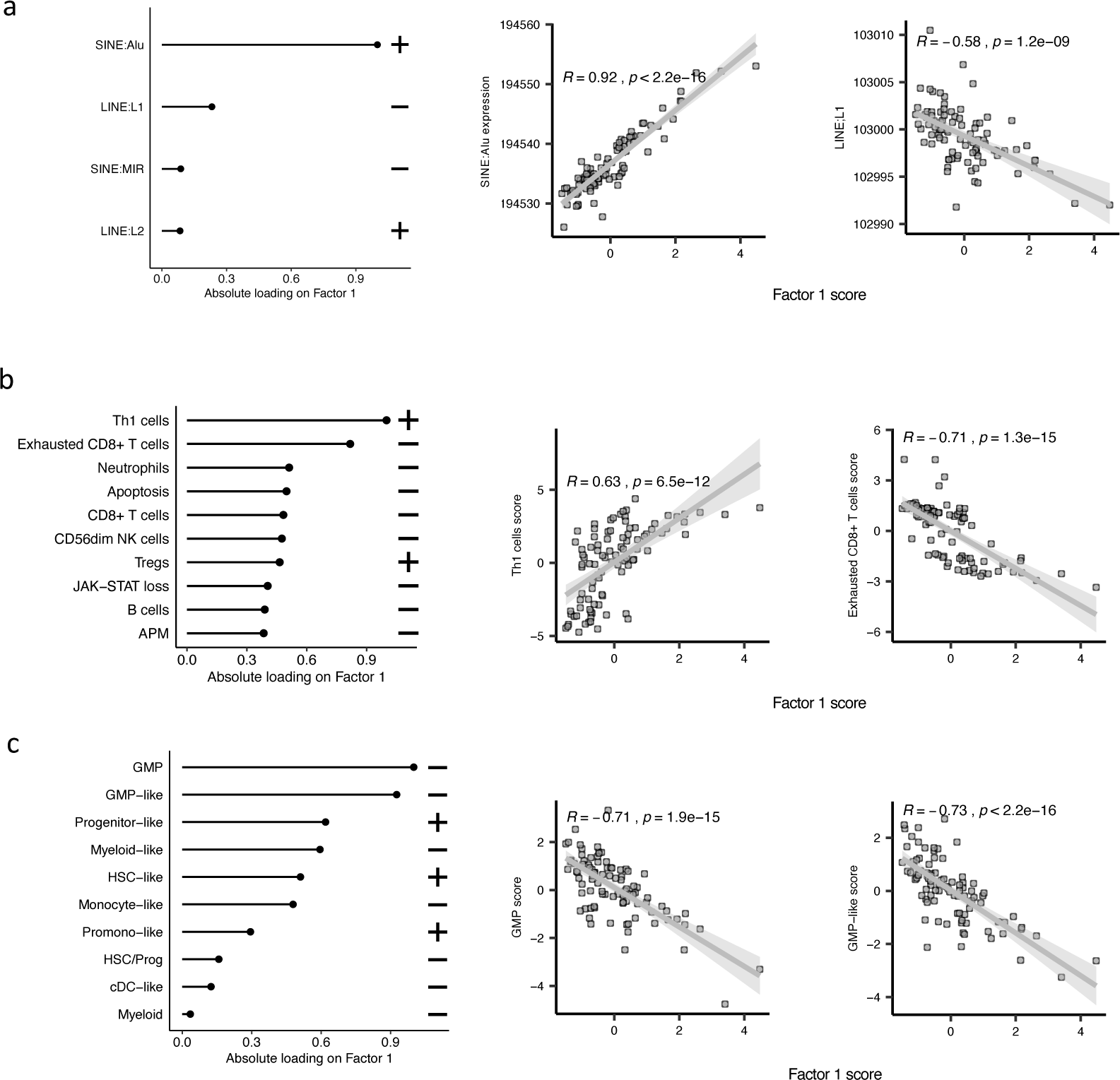
The top features influencing Factor 1 form various biological views in the BMMNC cohort. **a-c,** Absolute loadings of the top features and their positive or inverse correlations with Factor 1 from RTE expression, immune profile, and cell-type biological views, respectively. Pearson correlation coefficients (R) and p-values were displayed on top of the scatter plots.

**Supplementary Figure 2.**
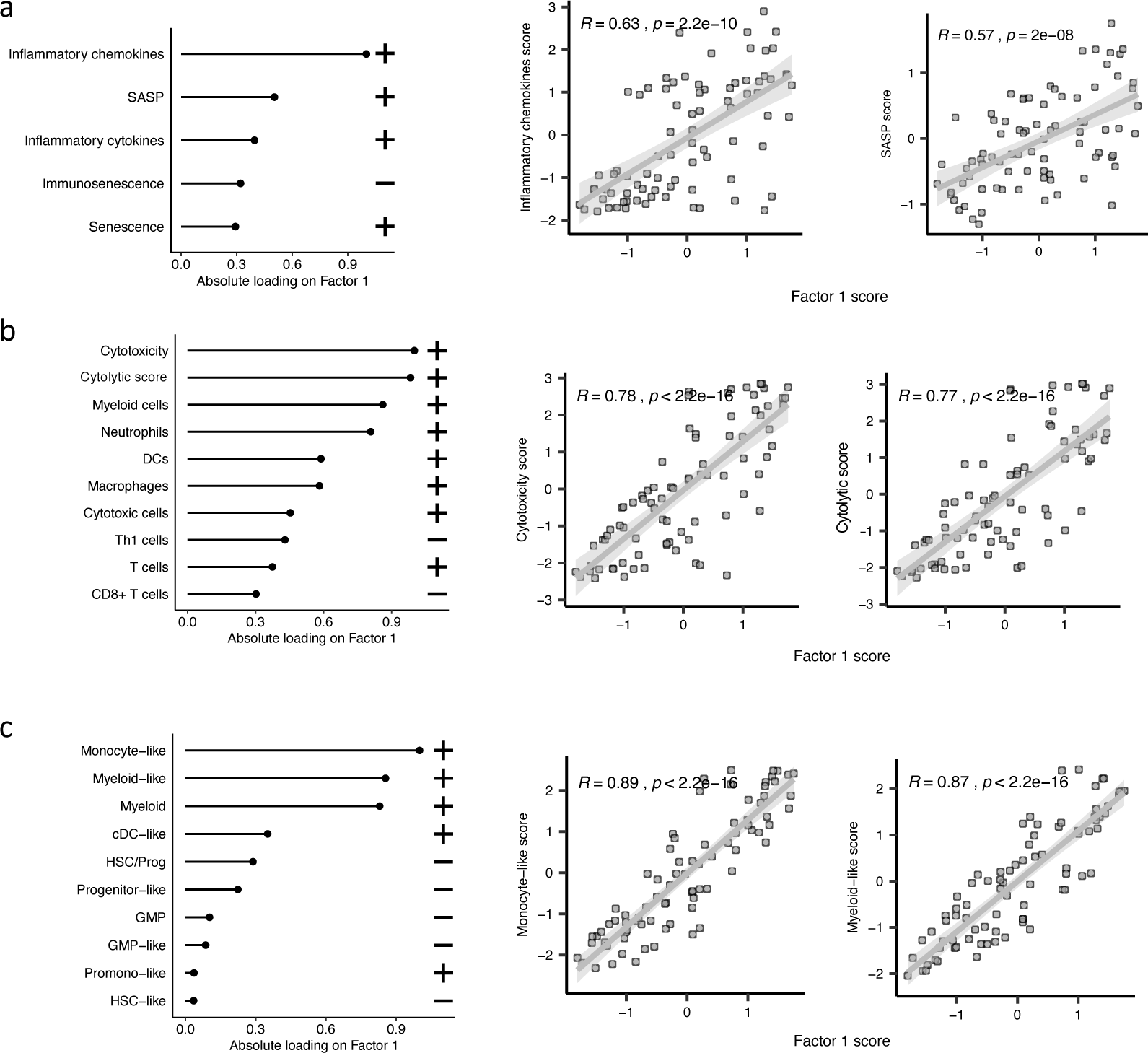
The top features influencing Factor 1 form various biological views in the CD34+ cohort. **a-c,** Absolute loadings of the top features and their positive or inverse correlations with Factor 1 from inflammation/aging, immune profile, and cell-type biological views, respectively. Pearson correlation coefficients (R) and p-values were displayed on top of the scatter plots.

**Supplementary Figure 3.**
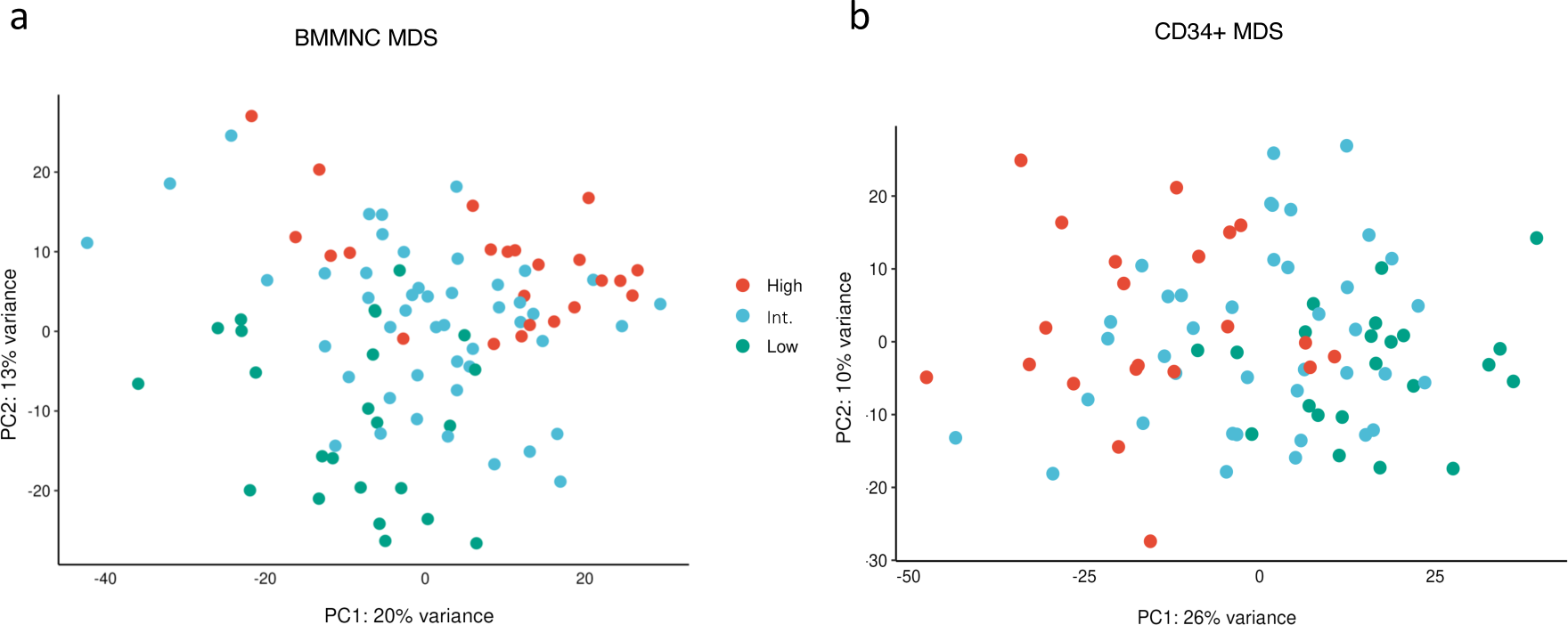
Factor 1 from the BMMNC and CD34+ MDS cohorts stratified the patients in the gene expression PCA plots. **a-b,** Principal component analysis on gene expression values of the BMMNC and CD34+ cohorts revealed separate clusters for patients split based on low (1^st^ quartile), intermediate (2^nd^ and 3^rd^ quartile), and high (4^th^ quartile) levels of Factor 1 in both cohorts. In both plots, the intermediate group is in between the high and low groups in the PCA plots.

**Supplementary Figure 4.**
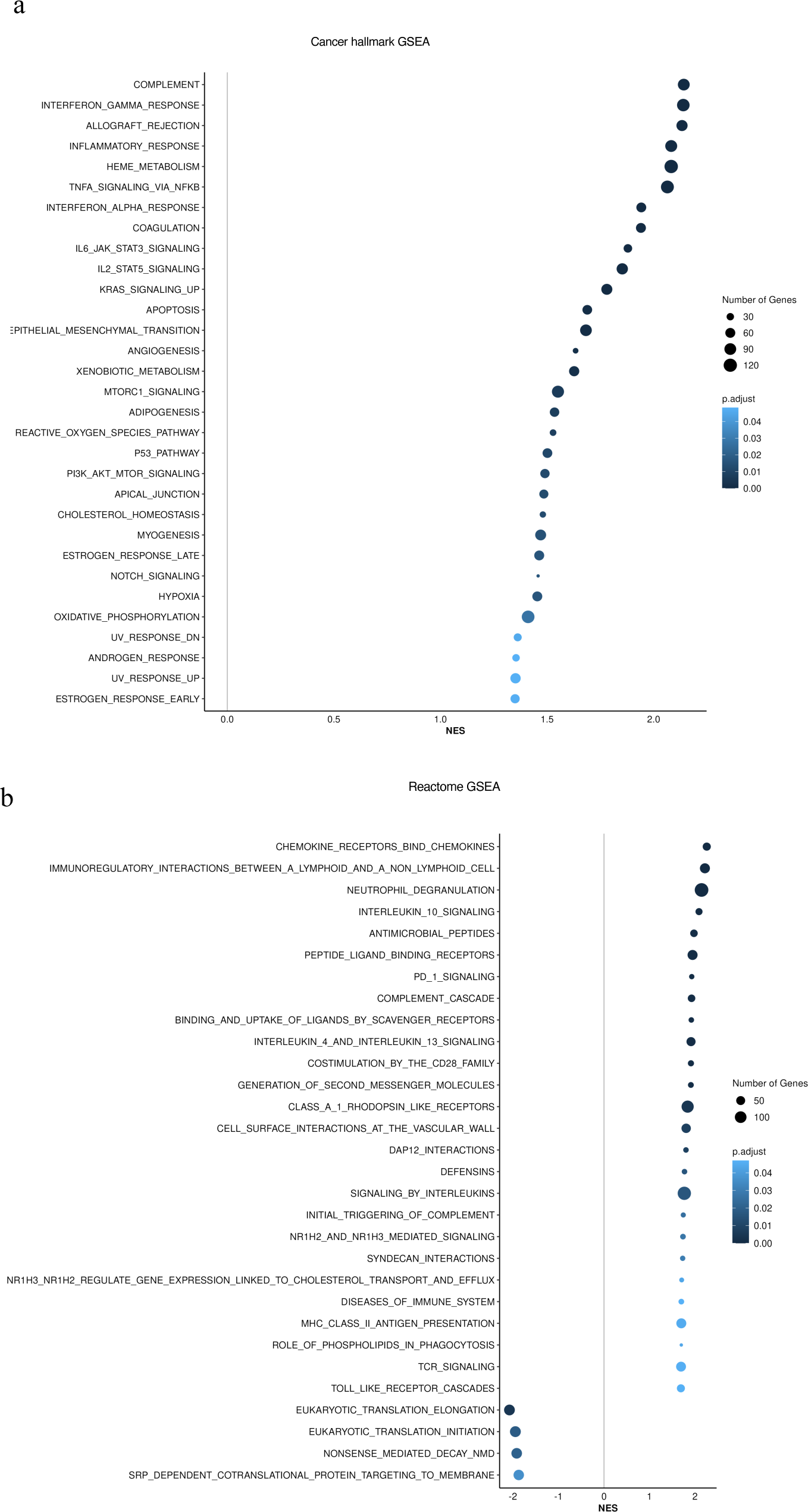
Gene set enrichment analysis (GSEA) identified upregulation of inflammation related pathways in patients having high (4th quartile) versus low (1st quartile) levels of Factor 1 in the CD34+ cohort. **a,** Upregulation of inflammatory, Interferons, TNFA, JAK-STAT signalling pathways within cancer hallmark gene sets. **b,** GSEA analysis using Reactome gene sets shows upregulation of gene sets associated with chemokines, Neutrophils deregulation and IL10 signalling.

**Supplementary Figure 5.**
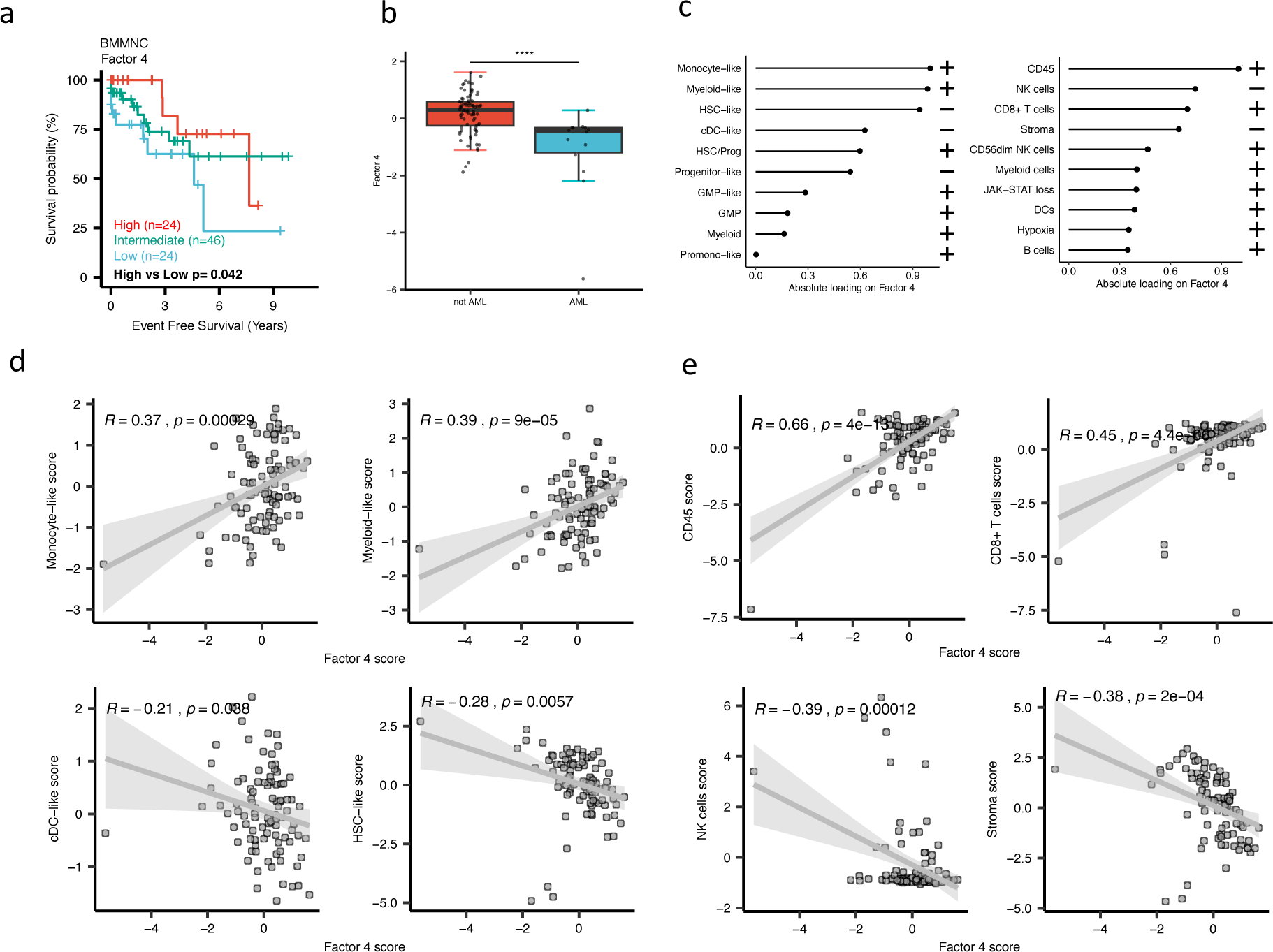
Characterisation of Factor 4 in the BMMNC cohort using gene signatures from multiple biological views. **a,** Kaplan-Meier plots for the BMMNC cohort where patients were split based on low (1^st^ quartile), intermediate (2^nd^ and 3^rd^ quartile), and high (4^th^ quartile) levels of Factor 4. The p-value was calculated using log-rank test on event-free survival values of high versus low groups. **b,** Boxplot depicting low level of Factor 4 for patients who progressed to AML. The significance was calculated with the Wilcox rank-sum test, and the significance was shown by **** (P < 0.0001). **c,** The absolute loading of the top features affecting Factor 4 in cell-type composition and immunology biological views in the BMMNC cohort. **d-e,** Positive or inverse correlations of selected top factors from the cell-type composition and immunology biological views with Factor 4. Pearson correlation coefficients (R) and p-values were displayed on top of the scatter plots.

**Supplementary Figure 6.**
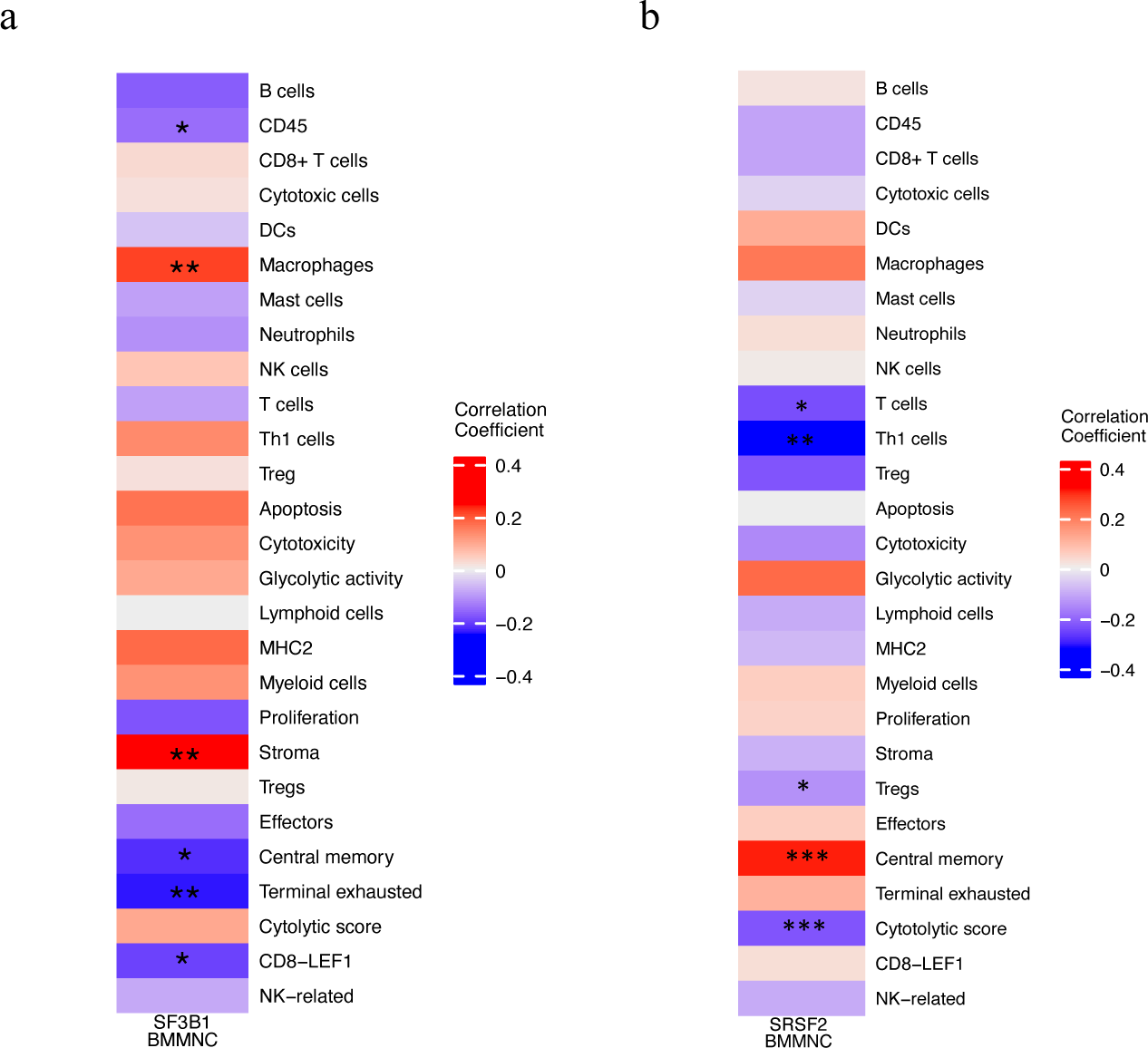
Correlation of selected immune profile gene sets with SF3B1 and SRSF2 mutations in the MDS BMMNC cohort. **a-b,** Correlation of a subset of immune profile features with SF3B1 and SRSF2 mutations in the BMMNC cohort, with red depicting a positive correlation and blue an inverse correlation with these mutations. The significances were calculated with the Wilcox rank-sum test, and the significant associations were shown by * (P < 0.05), ** (P < 0.01) or *** (P < 0.001).

**Supplementary Figure 7.**
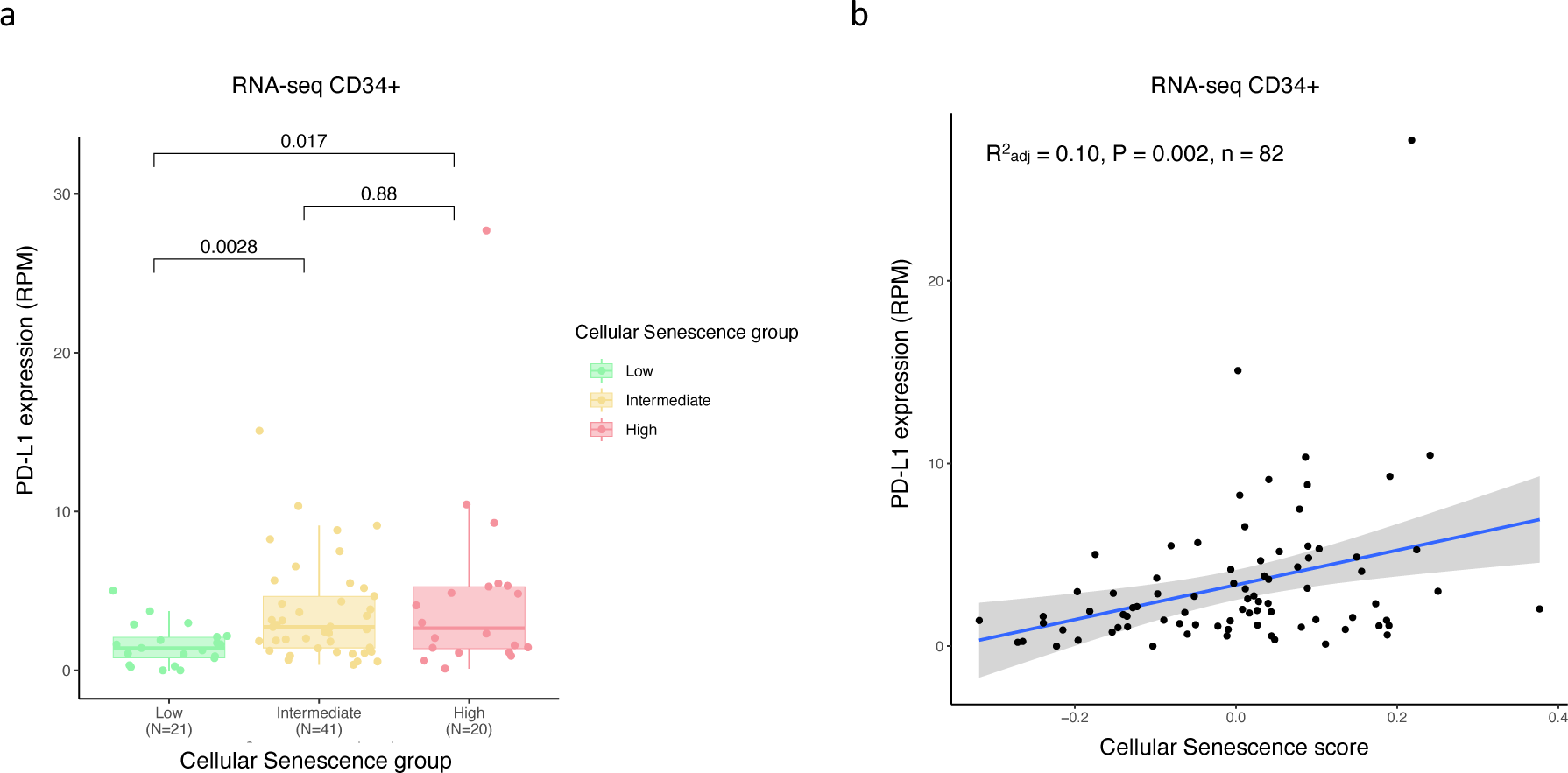
Positive correlation between PD-L1 expression and senescence score in BM CD34+ RNA-seq cohort. **a,** Boxplot showing higher expression of PD-L1 gene in patient groups with high (4^th^ quartile) and intermediate (2^nd^ and 3^rd^ quartile) vs low (1^st^ quartile) senescence levels. The significances were calculated with the Wilcox rank-sum. **b,** Scatter plot displaying a positive correlation between PD-L1 expression and senescence score.

